# Synergy-based Feedforward with Minimal Feedback Control Predicts Walking Over Multiple Cycles

**DOI:** 10.64898/2026.03.02.709098

**Authors:** Spencer T. Williams, Geng Li, Benjamin J. Fregly

**Affiliations:** Rice Computational Neuromechanics Lab, Department of Mechanical Engineering, Rice University, Houston, TX, United States

**Keywords:** Neuromusculoskeletal modeling, Predictive simulation, Model Personalization, Treatment Optimization, EMG-driven modeling, Muscle synergies, Neural feedback, Stroke

## Abstract

Neural feedback is important for the control of movement, and multiple neurological disorders (e.g., stroke, cerebral palsy, Parkinson’s disease, incomplete spinal cord injury) are characterized by altered neural feedback. Researchers have created numerous computational neuromusculoskeletal models controlled by simulated neural feedback mechanisms, but these models rarely represent actual human subjects and thus have not found practical clinical application. As a step toward designing patient-specific treatments for individuals with neurological disorders, this study used the Neuromusculoskeletal Modeling Pipeline to develop and evaluate a novel synergy-based feedforward (FF)+feedback (FB) control model using a personalized three-dimensional neuromusculoskeletal walking model of an actual human subject post-stroke. Experimental walking data collected from the subject were used to create the subject’s personalized walking model. Then for 5 calibration walking cycles, personalized synergy-based FF+FB control models were created. First, the personalized model was used to estimate lower body muscle activations consistent with the subject’s electromyographic, joint motion, and joint moment data. Second, five synergy activations per leg with associated synergy vectors were calculated that closely reconstructed the subject’s muscle activations and joint moments simultaneously. Third, nominal FF synergy activation controls were calculated by averaging the synergy activations for each leg. Fourth, the nominal FF synergy controls were scaled by 0, 25, 50, 75, 100, and 125%, and the gap in reproducing the subject’s muscle activations was filled by fitting FB synergy activation controls as a function of joint positions, velocities, and moments as surrogates for muscle lengths, muscle velocities, and tendon forces. Next for 3 testing walking cycles, the six synergy-based FF+FB models were used to control the subject’s personalized walking model in predictive simulations. The 100% FF model (which still had minimal FB) reproduced the testing walking cycles the most closely, and only the 75%, 100%, and 125% FF models predicted near-periodic walking motions using initial conditions consistent with experimental values. The 0, 25, and 50% FF models could generate near-periodic walking motions only when the initial conditions were allowed to diverge substantially from experimental values. Our findings suggest that predictive simulations of walking may require substantial feedforward control when modeling an actual human subject.

## 1 Introduction

Neural feedback is an essential component of the neuromuscular control of movement. Neural feedback helps individuals adapt their walking to new situations or perturbations (Severini and Zych, 2022), and reflexes help prevent dangerous slips and falls (McCrum et al., 2017). Disruptions to these mechanisms can produce muscle spasticity, a harmful velocity-dependent feedback mechanism that contributes to movement impairment and health hazards (Jansen et al., 2014; Trompetto et al., 2014; Ang et al., 2018; Falisse et al., 2018). Dysfunctional neural feedback is present in conditions such as cerebral palsy (Falisse et al., 2018), spinal cord injuries (Trompetto et al., 2014; Ang et al., 2018), stroke (Jansen et al., 2014; Ang et al., 2018), and Parkinson’s disease (Kim et al., 2009). However, a thorough understanding of how dysfunctional feedback mechanisms work remains elusive, hindering the development of more effective treatments for these conditions.

Researchers have explored the use of computational neuromusculoskeletal models to develop a better understanding of human reflex contributions to walking and movement impairments (Geyer and Herr, 2010; Song and Geyer, 2013; Jansen et al., 2014; Falisse et al., 2018; Haeufle et al., 2018; Di Russo et al., 2021; Lassmann et al., 2023; Su and Gutierrez-Farewik, 2023; Koseki et al., 2024). Reflex mechanisms are typically modeled at the level of individual muscles and have included stretch reflexes and Golgi tendon reflexes (Geyer and Herr, 2010; Jansen et al., 2014; Trompetto et al., 2014; Ang et al., 2018; Falisse et al., 2018; Haeufle et al., 2018). While complicated neural networks in the brain control movement, simple walking models can be successfully controlled by feedback mechanisms alone (Geyer and Herr, 2010). However, these models can be made more robust by combining feedforward and feedback control (Haeufle et al., 2018).

While most studies have assumed neural feedback mechanisms control individual muscles, several studies have explored feedback control of muscle synergies. In a computational study, neuromusculoskeletal models walked and ran at different speeds using similar control structures activating muscle synergies rather than individual muscles (Aoi et al., 2019). In an experimental study of subjects attempting to stand upright while being perturbed, muscle synergy controls could be fitted using feedback quantities related to a target motion, such as center of mass location (Safavynia and Ting, 2013), supporting the use of muscle synergies in human movement simulations. Additional studies have reported that neural feedback for the control of movement in general and walking in particular may in part come through muscle synergies (Ting and McKay, 2007; Chvatal and Ting, 2013). It is notable, however, that most past simulation studies have predicted general movement characteristics or limited experimental data rather than the response of a specific subject under new conditions. Furthermore, no previous study has addressed the issue of how much feedback control is needed to simulate walking performed by an actual human subject.

As a step toward addressing this question, this study describes the development and evaluation of a novel process for creating personalized synergy-based feedforward (FF)+feedback (FB) neural control models to predict three-dimensional (3D) walking for an individual post-stroke. The study is made possible by a personalized neuromusculoskeletal walking model created from experimental walking data collected from the subject. Personalized FF+FB neural control models are implemented using muscle synergies. Models with different ratios of FF to FB control, combined using an approach from a previously published study (Haeufle et al., 2018), are calibrated using 5 cycles of walking data and tested in predictive simulations using 3 additional cycles of walking data withheld from calibration. The relative contributions of FF and FB control are explored with the ultimate goal of facilitating the design of personalized treatments that target impairments in a individual’s control mechanisms. Personalized neural feedback models similar to the ones investigated here may eventually help researchers and clinicians identify optimal treatment plans for individuals with movement impairments caused by various neurological disorders.

## 2 Materials and Methods

### 2.1 Experimental Data

For this modeling study, we used published experimental walking data collected with written informed consent from a male stroke survivor (age 79 years, LE Fugl-Meyer Motor Assessment 32/34 pts, right-sided hemiparesis, height 1.7 m, mass 80.5 kg) (Meyer et al., 2016). The walking data included marker-based motion capture data collected using a modified Cleveland Clinic full-body marker set with additional foot markers (Reinbolt et al., 2005) (Vicon Corp., Oxford, UK), ground reaction force and moment data collected using a split-belt instrumented treadmill (Bertec Corp., Columbus, OH, USA), and electromyographic (EMG) data collected from 16 muscles per leg (Motion Lab Systems, Baton Rouge, LA, USA). Data collected during a standing static pose were used for model scaling purposes (see below), while data collected during 30 seconds of treadmill walking at a self-selected speed of 0.5 m/s were used for the feedback control study.

EMG data for each leg were collected using a combination of surface and fine wire electrodes. Surface electrodes were used for gluteus maximus and medius, semimembranosus, biceps femoris long head, rectus femoris, vastus medialis and lateralis, medial gastrocnemius, tibialis anterior, peroneus longus, and soleus. Fine wire electrodes were used for adductor longus, iliopsoas, tibialis posterior, extensor digitorum longus, and flexor digitorum longus. Raw EMG data were converted to muscle excitation envelopes using standard EMG processing methods (Lloyd and Besier, 2003).

### 2.2 Neuromusculoskeletal Modeling

With the exception of fitting FB models, all modeling and simulation tasks in this study were performed using OpenSim (Seth et al., 2018) and the Neuromusculoskeletal Modeling (NMSM) Pipeline (Hammond et al., 2025). Code, input data, XML settings files, and output data are available on SimTK.org at the link provided in the Data Availability Statement. All NMSM Pipeline tools perform numerical optimizations either to personalize some aspect of a subject’s neuromusculoskeletal model or to predict the subject’s movement under some new condition. We encourage readers interested in how NMSM Pipeline tools work to review the published NMSM Pipeline article (Hammond et al., 2025). This manuscript includes a flowchart of how these tools and their outputs interact (Figure 1).

**Figure 1.**
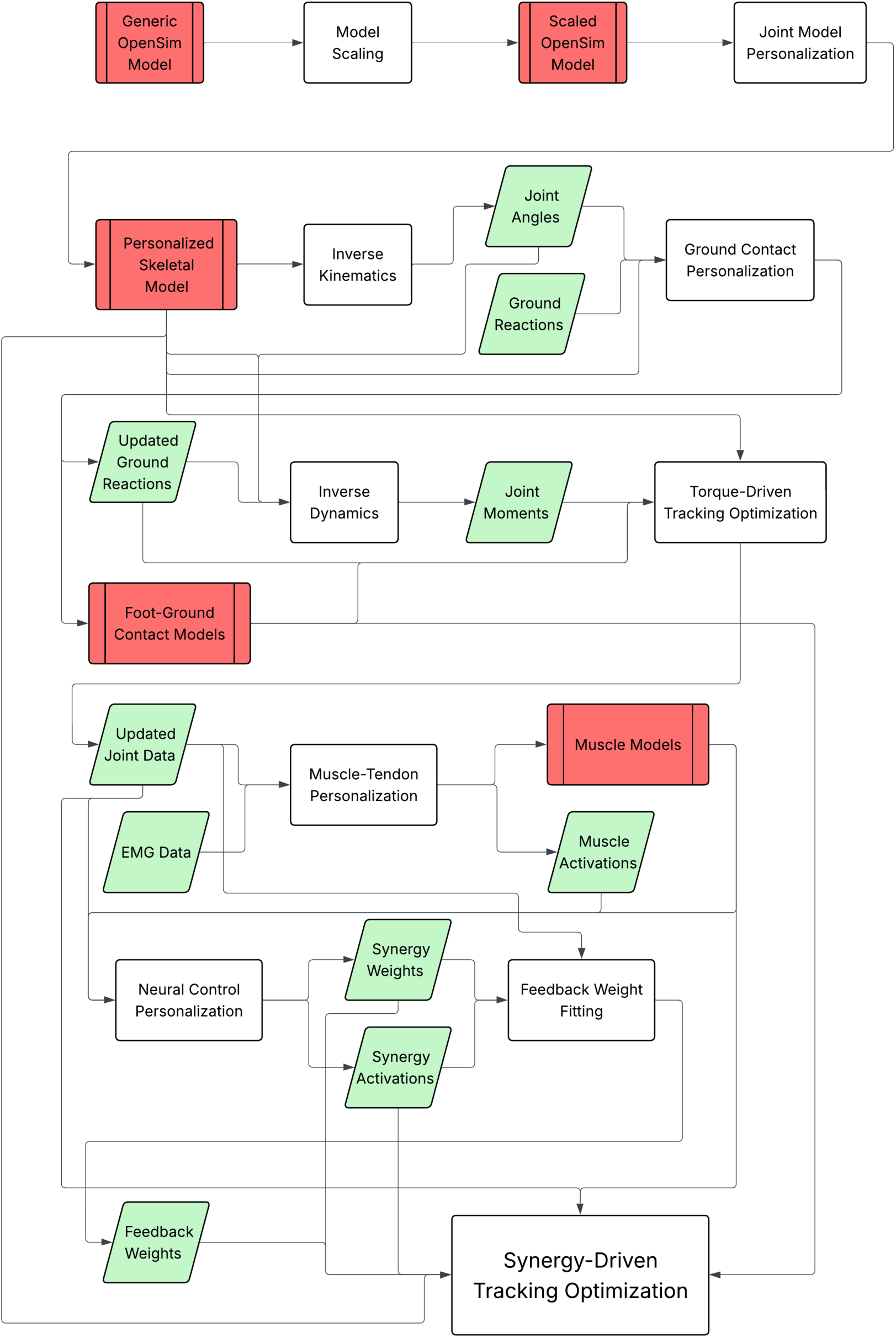
A flowchart outlining the high-level processes used in our Methods.

In all NMSM Pipeline tools, optimization cost function terms are calculated using the following equation:

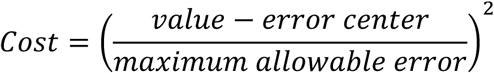

This equation is designed to penalize value deviations from error centers, which are defined using either nominal values for model parameters (e.g., an optimal muscle fiber length) or time-varying experimental data (e.g., joint angles). Values that deviate by more than the user-specified “maximum allowable” error create a normalized error greater than one, which becomes even larger when squared. Optimization problem formulations and non-default settings are described in detail in Supplementary Material Tables 1 through 7.

#### 2.2.1 Joint and Ground Contact Model Personalization

A generic full-body OpenSim (Seth et al., 2018) musculoskeletal model (Hammond et al., 2025) derived from a previously published OpenSim model (Rajagopal et al., 2016) was used as a baseline. Forearm pronation-supination and knee adduction degrees of freedom (DOFs) were locked, leaving the generic model with 31 DOFs. These DOFs included a 6 DOF ground-to-pelvis joint, two 3 DOF hip joints, two 1 DOF knee joints, two 1 DOF ankle joints, two 1 DOF subtalar joints, two 1 DOF toe joints linking the hindfoot and toes, a 3 DOF back joint, two 3 DOF shoulder joints, and two 1 DOF elbow joints. The model included 86 Hill-type muscle-tendon actuators with rigid tendons (43 per leg). The model was scaled in OpenSim using the static pose data with bodies and joints scaled symmetrically.

The skeletal geometry and joint parameters of the scaled model were personalized to the subject’s experimental data using the Joint Model Personalization tool within the NMSM Pipeline (Hammond et al., 2025). The tool optimized joint axis orientations, isotropic body scale factors, and marker locations to reduce inverse kinematics (IK) marker distance errors while seeking to reproduce the motion of a portion of the 30 second walking trial. The tool was allowed to change joint axis orientation parameters in the knee, ankle, and subtalar joints, scaling of the pelvis, femur, and tibia bodies, and the locations of markers attached to the femur and tibia bodies. The cost function penalized IK marker distance errors. The calibrated model was then used with OpenSim’s Inverse Kinematics tool to solve for joint angles that reproduced the experimental marker motion data as closely as possible. The IK results were filtered using a fourth-order zero phase lag Butterworth filter with a cutoff frequency of 7/tf Hz, where tf is the average period of a gait cycle from the 30 second recording (Hug, 2011).

After the skeletal structure was personalized and IK joint angles were calculated, the Ground Contact Model Personalization (GCP) tool within the NMSM Pipeline (Hammond et al., 2025) was used to generate a subject-specific elastic foundation foot-ground contact model that was the same for both feet. Prior to running the tool (and for future steps), the experimental ground reaction forces and moments were filtered using a second-order zero phase lag Butterworth filter with a cutoff frequency of 7/tf Hz. The GCP tool automatically placed 37 viscoelastic+friction contact elements across the hindfoot and toes segments of each foot and calibrated each contact element’s stiffness coefficient, a common nonlinear damping coefficient, a common dynamic friction coefficient, and a common spring resting length defining the height at which the contact elements penetrated the ground. To calibrate these parameters, the optimization cost function penalized errors in reproducing all three components of the experimental ground reaction forces and moments while making minimal adjustments to the experimental foot kinematics with respect to ground. To prevent unrealistic discontinuities in contact element stiffness values, the cost function also minimized the deviation of each contact element stiffness value away from neighboring values using a Gaussian-weighted average, where stiffness values from nearer elements had a higher weight. In addition, to make the ground reaction data more consistent with the marker motion data and the dynamics of the skeletal model, the cost function made small adjustments to the location of each force plate in the lab coordinate system (Williams et al., 2025). The adjusted ground reaction data were used when calculating joint moments with OpenSim’s Inverse Dynamics (ID) tool.

#### 2.2.2 Torque-Driven Tracking Optimization

Joint angle, ID, and ground reaction data were used in the Tracking Optimization (TO) tool within the NMSM Pipeline (Hammond et al., 2025) to generate dynamically consistent torque-controlled walking simulations of individual gait cycles to be used in place of experimental data. From the full 30 seconds of walking data, 8 gait cycles with clean marker motion, ground reaction, and EMG data were identified. Each gait cycle started with right foot heelstrike, and IK, ID, and ground reaction data for these 8 cycles were split into separate data files, each splined to 101 time points. The data for each gait cycle were used to provide an initial guess and tracked quantities for direct collocation optimal control motion prediction problems solved using the TO tool. The values of coordinate positions, velocities, and accelerations were allowed to change, resulting in different ground reactions and joint loads, and torque controllers were used to control the motion of the hip, knee, ankle, and subtalar joints. The arms and torso did not have associated controllers, but their coordinate positions, velocities, and accelerations were included as design variables. The cost function minimized tracking errors in joint angles, joint velocities, joint accelerations, ground reaction forces and moments, ID joint loads, and heel marker positions and velocities. Constraints were used to maintain joint coordinate and joint speed periodicity and initial and final values of ground reaction forces and moments. Additional constraints enforced that torque controls calculated during the optimization stay within 0.1 Nm of their respective ID joint loads and that residual ID loads calculated for the pelvis, corresponding to the 6 DOF ground-to-pelvis joint, were less than 1 N or 0.1 Nm in magnitude. Dynamically consistent TO results for each of the 8 gait cycles were used in future steps.

#### 2.2.3 Muscle-tendon and Neural Control Model Personalization

The Muscle Tendon Personalization (MTP) tool within the NMSM Pipeline (Hammond et al., 2025) was used to calibrate muscle-tendon model properties. Based on the TO results for each of the 8 gait cycles, time-varying muscle length, velocity, and moment arm data were calculated for each muscle. These data were filtered with using the same protocol as IK joint angles. EMG data were high-pass filtered at 40 Hz, demeaned, rectified, low-pass filtered with a cut-off frequency of 3.5/tf Hz (Meyer et al., 2016), normalized to a maximum value of 1, split into separate data files for each cycle, and splined to 101 time points plus a cycle-dependent number of padding frames. The padding frames added 200 ms of data before and after the one-cycle time range used by ID to accommodate EMG electromechanical delay. The MTP tool calibrated muscle-tendon parameters and solved for time-varying activations for each muscle using EMG data. Muscle-specific values for electromechanical delays, activation time constants, activation nonlinearity constants, EMG scale factors, optimal muscle fiber lengths, and tendon slack lengths were calculated. In addition, synergy extrapolation (Ao et al., 2022) was used to estimate the excitations of hip rotator muscles, tensor fasciae latae, and sartorius, which muscles that did not have experimental EMG data. The cost function penalized: 1) errors in hip flexion, hip adduction, knee, ankle, and subtalar joint moments produced by muscles relative to ID loads from the torque-driven TO simulations; 2) extrapolated muscle excitations; 3) changes to measured excitations made by the synergy extrapolation process; 4) deviations from error centers in activation time constants, optimal muscle fiber lengths, tendon slack lengths, EMG scale factors, and normalized muscle fiber lengths; 5) activation nonlinearity constants; 6) passive muscle forces; 7) muscle excitations; 8) deviations within defined muscle groups in normalized fiber length, and 8) electromechanical delays. The optimization was performed for all 8 gait cycles simultaneously to ensure all cycles used consistent muscle-tendon properties.

Next, the Neural Control Model Personalization (NCP) tool within the NMSM Pipeline (Hammond et al., 2025) was used to impose a synergy structure on the MTP muscle activations while ensuring consistency with ID joint moments. From the 8 cycles of TO results, the 5 cycles with the largest deviations of IK, ID, and ground reaction data from their mean curves were selected for fitting a neural back model, thereby capturing the most variation available during the calibration process. The remaining 3 cycles were set aside for testing feedback models. Using the 5 fitting cycles, NCP solved for 5 sets of synergy activations unique to each leg and one set of synergy vector weights common to both legs. Use of common synergy vectors between paretic and non-paretic legs enabled a direct comparison of synergy activation levels between the two legs and is supported by published studies (Cheung et al., 2009; Clark et al., 2010; Komaris et al., 2025). Consequently, asymmetry in the subject’s control was captured by his synergy activations. The number of synergies per leg was chosen by comparing NCP results for different numbers of synergies, where the goal was to achieve a total variability accounted for (VAF) > 90% while avoiding any highly similar pairs of synergy vectors with a cosine similarity > 0.7. While the 6-synergy solution had a total VAF just above 90%, it also possessed a pair of highly similar synergy vectors with a cosine similarity above 0.7. Consequently, the 5-synergy solution was chosen, which possessed a total VAF of 88% and maximum cosine similarity of 0.63 for any pair of synergy vectors. The cost function penalized errors in tracking MTP muscle activations and TO joint moments as well as activation deviations within specified muscle groups. Synergy activations were calculated for each cycle using a single set of synergy vectors for all cycles and both legs. The mean synergy activations for each leg were used as nominal feedforward synergy activations in future steps.

Finally, FB synergy activations were fitted as functions of joint angles, joint velocities, and ID moments, which were used as surrogates for muscle lengths, muscle velocities, and tendon forces typically used when modeling muscle stretch reflexes and Golgi tendon reflexes. For each assumed percentage of nominal FF control (0%, 25%, 50%, 75%, 100%, and 125%), the gap between each total synergy activation and each corresponding scaled nominal FF synergy activation was calculated for each gait cycle, and a synergy activation linear feedback model was fitted to fill these gaps. For each synergy activation gap *s* (101 x 1, where 101 is the number of time frames for one gait cycle), a linear system of equations was formulated that related joint angles *q* (hip flexion and adduction, knee, ankle, and subtalar angles), joint velocities *q̇*, and joint moments *M* to the synergy activation gap via unknown feedback gains *K*_1_ (1 x 1), *K_q_* (5 x 1, where 5 is the number of controlled joints in each leg), *K*_q_ (5 x 1), and *K*_%_ (5 x 1):

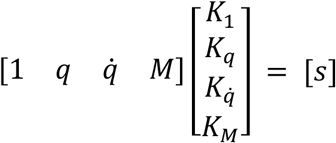

Stacking data for 5 gait cycles produces a linear system of 505 equations in 16 unknowns. Though this linear system appears to be overdetermined, in practice it was poorly conditioned, necessitating a solution approach that sought to satisfy the linear system of equations above while also minimizing the magnitude of the feedback gains. This approach was implemented in MATLAB using the non-linear least squares optimization algorithm *lsqnonlin*. The quality of fit for the synergy activation FB models was assessed using R^2^ values comparing nominal FF+calculated FB synergy activations with synergy activations produced by the NCP tool.

#### 2.2.4 Synergy-Driven Predictive Simulations with Neural Feedback

Each of the 6 synergy-based FF+FB models was used in a synergy-driven predictive simulation performed with the TO tool to evaluate how well the 3 testing gait cycles withheld from calibration could be reproduced. As in the torque-driven simulations, the arms and torso did not have associated controllers, but their coordinate positions, velocities, and accelerations were included as design variables. The cost function penalized errors in upper-body, hip internal-external rotation, and toes joint angles and velocities along with torso orientation and angular velocity with respect to ground. These cost terms accounted for portions of the model that were not controlled by muscles. Nominal FF synergy activations were included in the optimal control problem as controls and constrained to have changes of less than 0.01, thereby enforcing their shapes without negatively impacting the performance of the optimization. Additional constraints were used to ensure that initial values of joint angles, joint velocities, and ground reactions remained close to experimental initial values (within ± 2 deg, 20 deg/s, 2 cm, 2 cm/s, 10 N, and 2 Nm), the predicted motion was near-periodic, dynamic consistency was maintained by keeping muscle-produced joint moments within 1 Nm of their corresponding ID joint loads, and residual ID loads applied to the pelvis were less than 1 N or 0.1 Nm in magnitude. To improve computational speed, we also fitted surrogate geometric models of muscle-tendon lengths, velocities, and moment arms using Latin hypercube sampling around joint angle trajectories obtained from one representative walking cycle. The accuracy of predicted joint angles, joint moments, and ground reactions was assessed using root mean square error (RMSE) relative to the torque-driven TO results.

The success of each predictive simulation was categorized in a binary fashion. If a predictive simulation produced a near-periodic walking motion while remaining within the specified initial condition constraints, the predictive simulation was categorized as “successful.” If a predictive simulation could not satisfy these requirements, it was categorized as “unsuccessful” and was then re-run with the constraints on initial joint angles, joint velocities, and ground reactions removed.

## 3 Results

The synergy control structure produced by NCP corresponded well to known information about the subject’s control of muscles. Synergy vectors calibrated using the 5 calibration gait cycles exhibited visibly distinct distributions between synergies (Figure 2). As synergy vectors were symmetric between legs, the subject’s asymmetric control was manifested through his synergy activations in both shape and magnitude (Figure 3). The total variability accounted for (VAF) of muscle activations within individual gait cycles was 88.46 ± 1.05%. To compare calibration and testing cycles, we calculated how close the muscle activations for the testing cycles were to the mean muscle activations for the calibration cycles. The testing cycle mean muscle activations fell within 1.75 standard deviations of the mean muscle activations for the calibration cycles.

**Figure 2.**
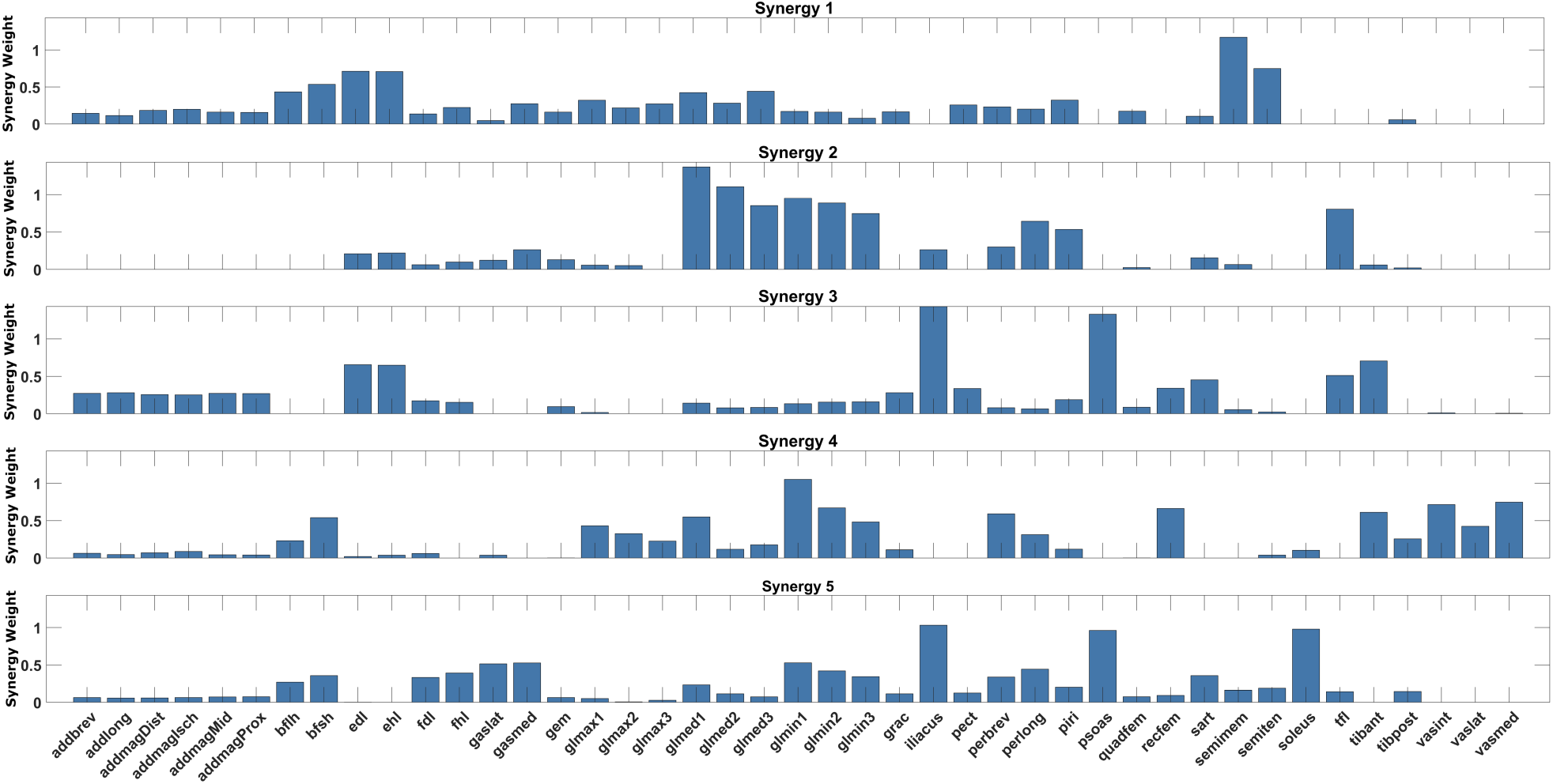
Optimized synergy weights derived from 5 fitting gait cycles.

**Figure 3.**
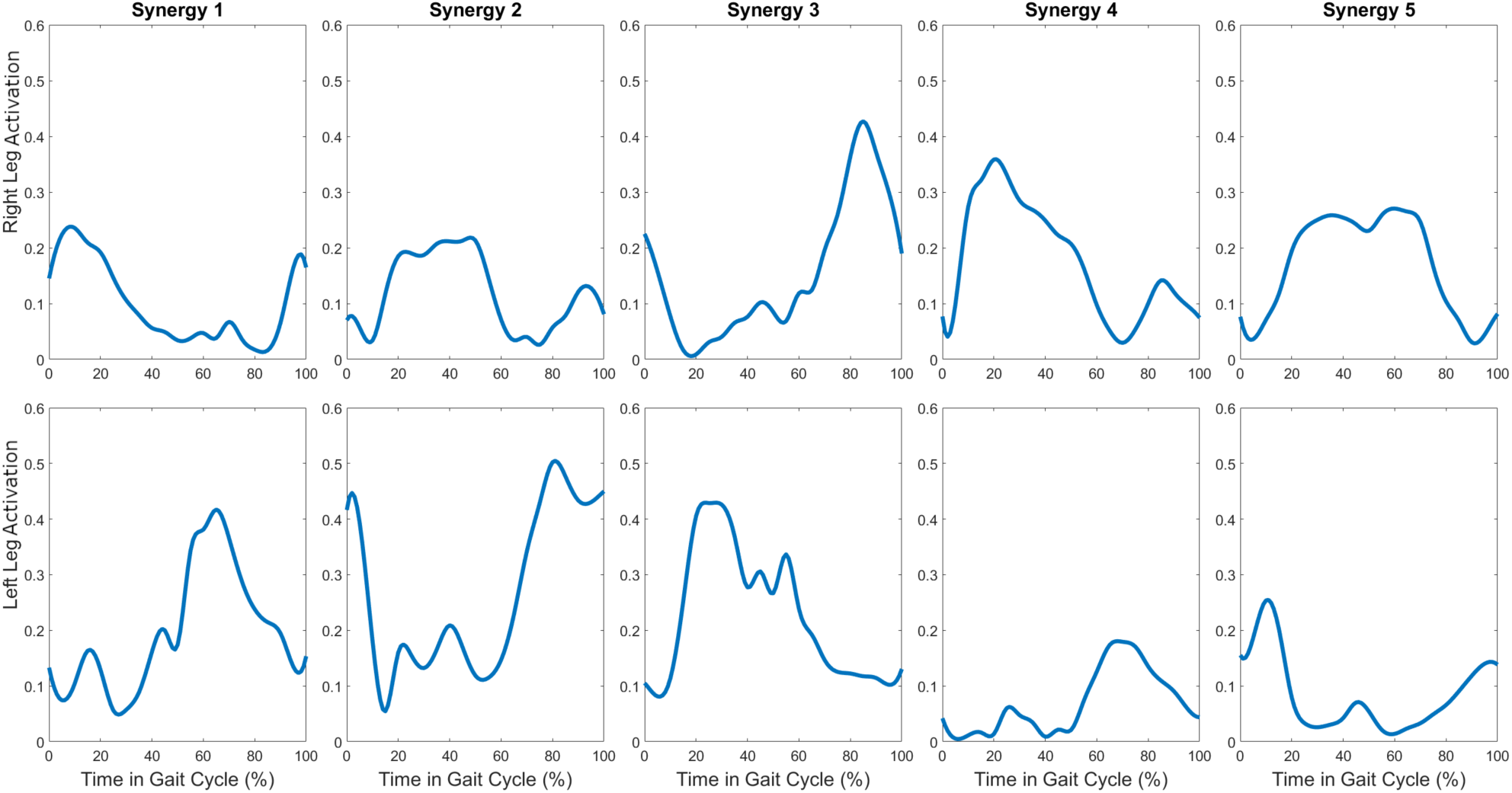
Mean synergy activations from 5 fitting gait cycles used as feedforward activations.

Calibrated synergy activation FB gains closely reproduced calculated synergy activation gaps. R^2^ values for fitting all 10 synergy activations (5 per leg) in all 6 feedforward magnitude cases were ≥ 0.9 with the exception of a single synergy in the 125% feedforward case (Table 1). Visual inspection of the fitted total synergy activations was consistent with this high accuracy (Figure 4).

**Figure 4.**
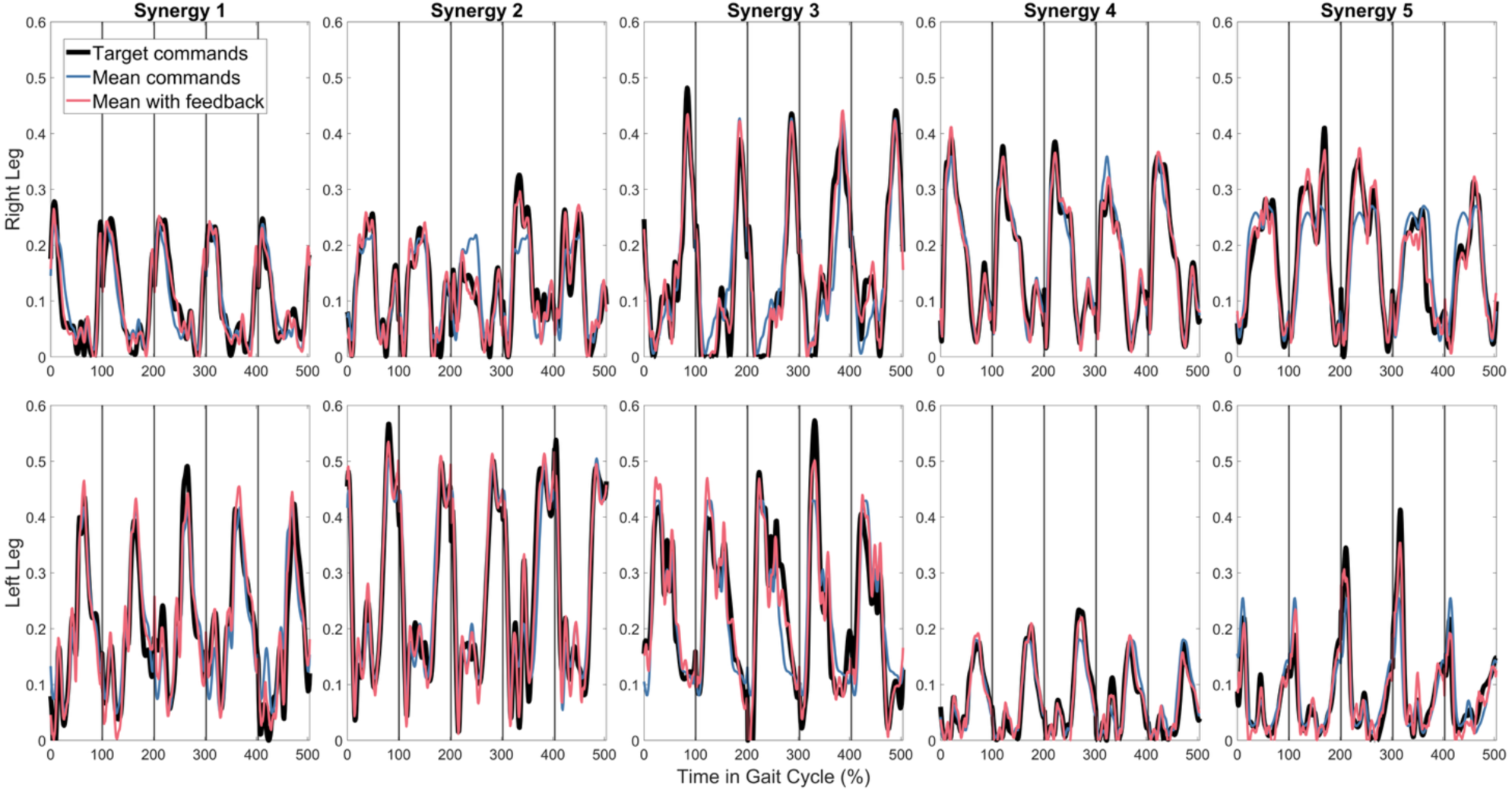
Fit of reference synergy activations (black) by feedback model at 100% feedforward contribution (red) compared to mean synergy activations (blue).

**Table 1.** R^2^ values for fitting synergy activations at each FF contribution level.

| FF contribution | 0% | 25% | 50% | 75% | 100% | 125% |
| --- | --- | --- | --- | --- | --- | --- |
| Synergy 1 (R) | 0.978 | 0.976 | 0.973 | 0.959 | 0.950 | 0.939 |
| Synergy 2 (R) | 0.981 | 0.978 | 0.973 | 0.943 | 0.928 | 0.908 |
| Synergy 3 (R) | 0.975 | 0.976 | 0.975 | 0.967 | 0.963 | 0.957 |
| Synergy 4 (R) | 0.981 | 0.986 | 0.987 | 0.982 | 0.979 | 0.971 |
| Synergy 5 (R) | 0.977 | 0.979 | 0.980 | 0.958 | 0.953 | 0.946 |
| Synergy 1 (L) | 0.960 | 0.959 | 0.954 | 0.931 | 0.917 | 0.898 |
| Synergy 2 (L) | 0.992 | 0.991 | 0.989 | 0.977 | 0.967 | 0.953 |
| Synergy 3 (L) | 0.947 | 0.945 | 0.941 | 0.927 | 0.915 | 0.900 |
| Synergy 4 (L) | 0.981 | 0.981 | 0.979 | 0.961 | 0.948 | 0.930 |
| Synergy 5 (L) | 0.959 | 0.960 | 0.959 | 0.931 | 0.919 | 0.903 |

For the predictive simulations using synergy-based FF+FB control, only the 75%, 100%, and 125% FF cases were successful. The most accurate case was 100% FF control with FB, which predicted joint angles (Figure 5) and joint moments (Figure 6) that were visually similar to the reference data. While the 0%, 25%, and 50% FF cases were unsuccessful, they still produced near-periodic walking motions once the constraints were removed on remaining close to experimental initial conditions for joint positions, joint velocities, and ground reactions. However, the predicted walking motions were visually different from the reference walking motions (Figure 7), with the model taking wider, shorter strides that increased the base of support compared to the experimental motions. These visual large deviations were consistent with larger errors in predicted joint angles (Table 2), joint velocities (Table 3), and joint moments (Table 4) compared to successful simulations. For all cases, predictive simulations for the three testing gait cycles achieved similar levels of accuracy (Tables 2 through 4, see Supplementary Tables 12 through 20 for individual cycles) and were all either successful or unsuccessful when the constraints on remaining close to experimental initial conditions were enforced. These many variations were possible as Tracking Optimization problems converged within a few hours (see Supplementary Table 8).

**Figure 5.**
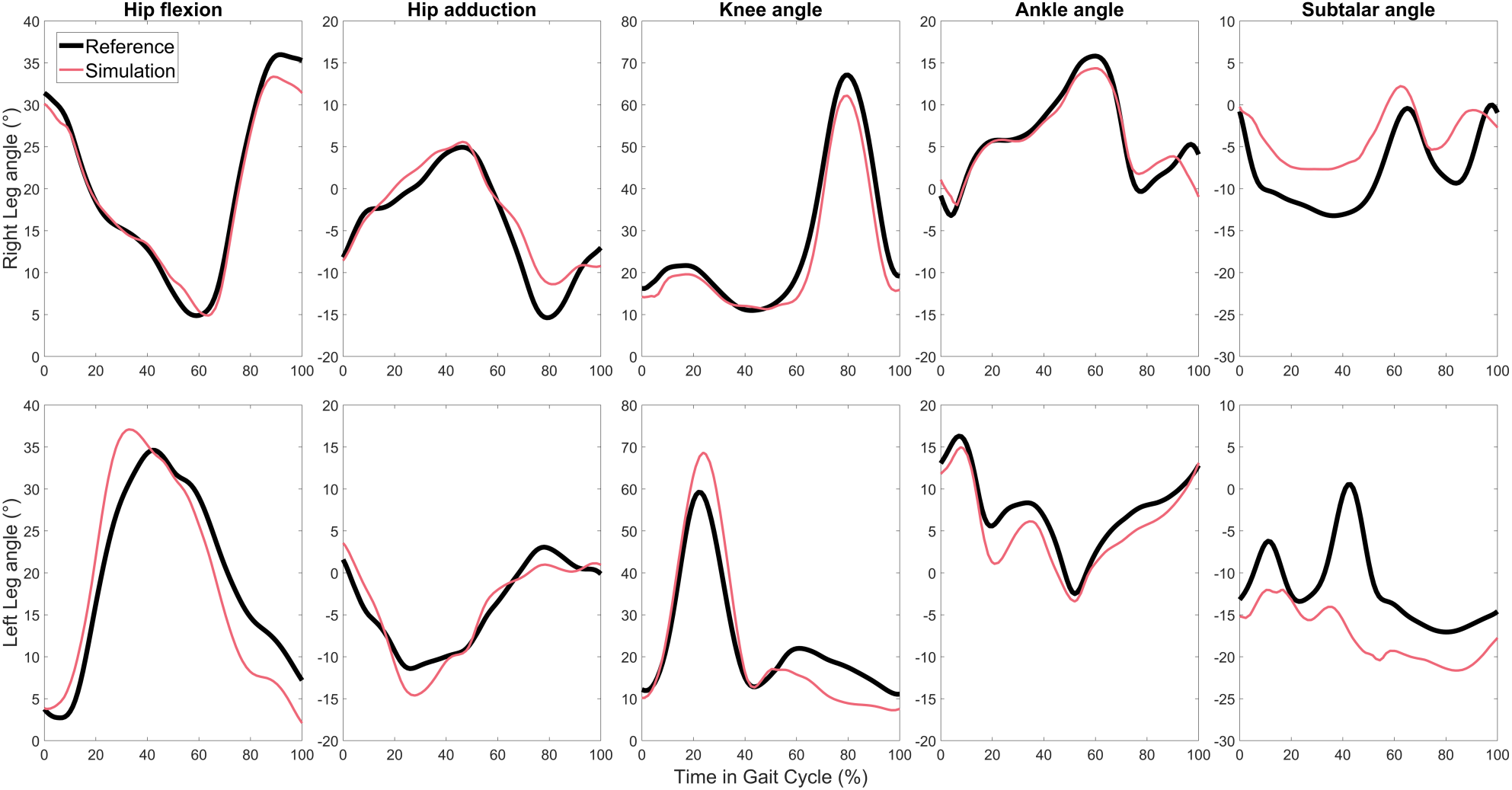
Reference (black) and predicted (red) joint angles using the feedback model with 100% feedforward contribution for testing cycle 3, which was the most successful cycle overall.

**Figure 6.**
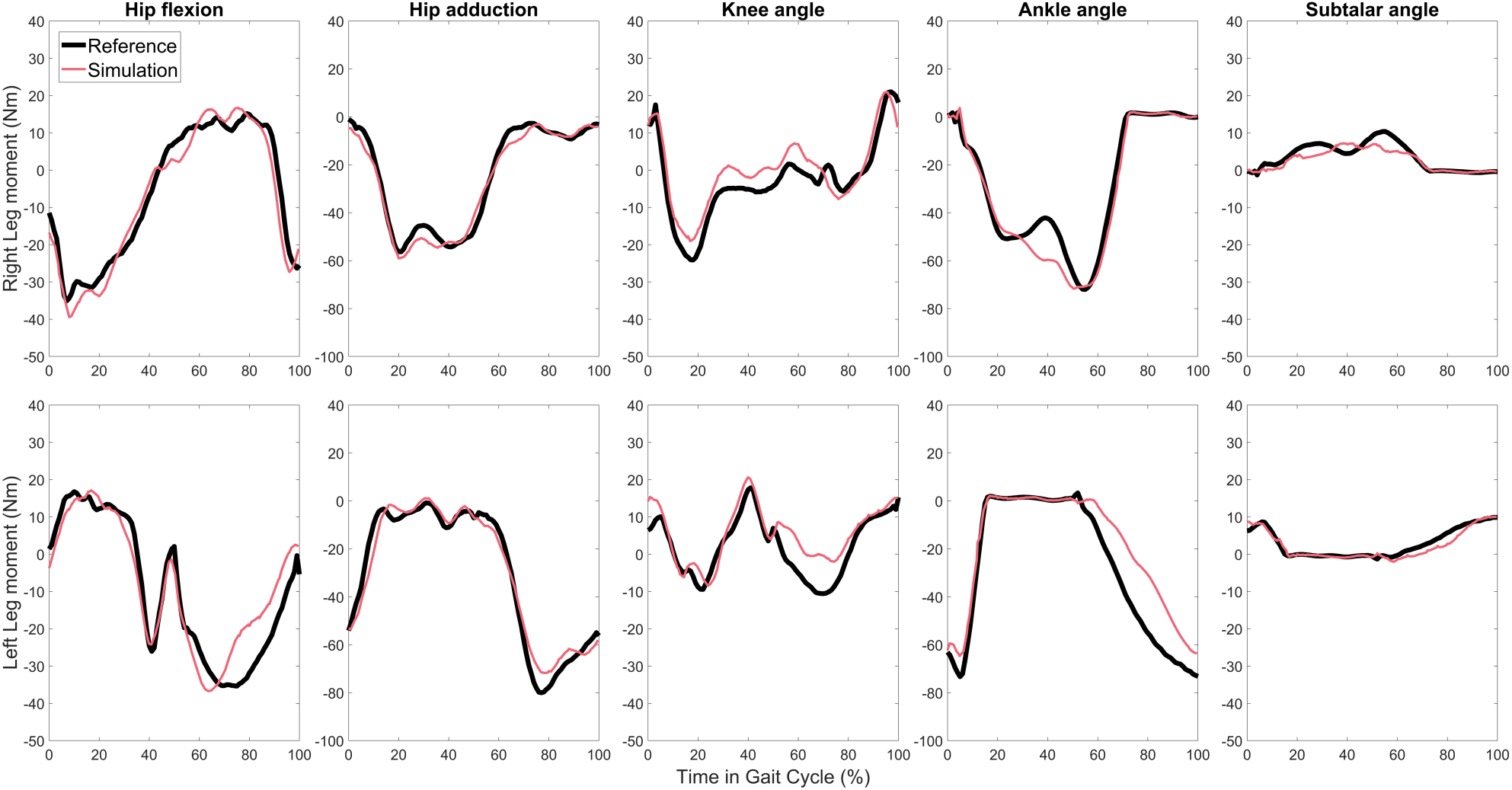
Reference (black) and predicted (red) joint moments using the feedback model with 100% feedforward contribution for testing cycle 3.

**Figure 7.**
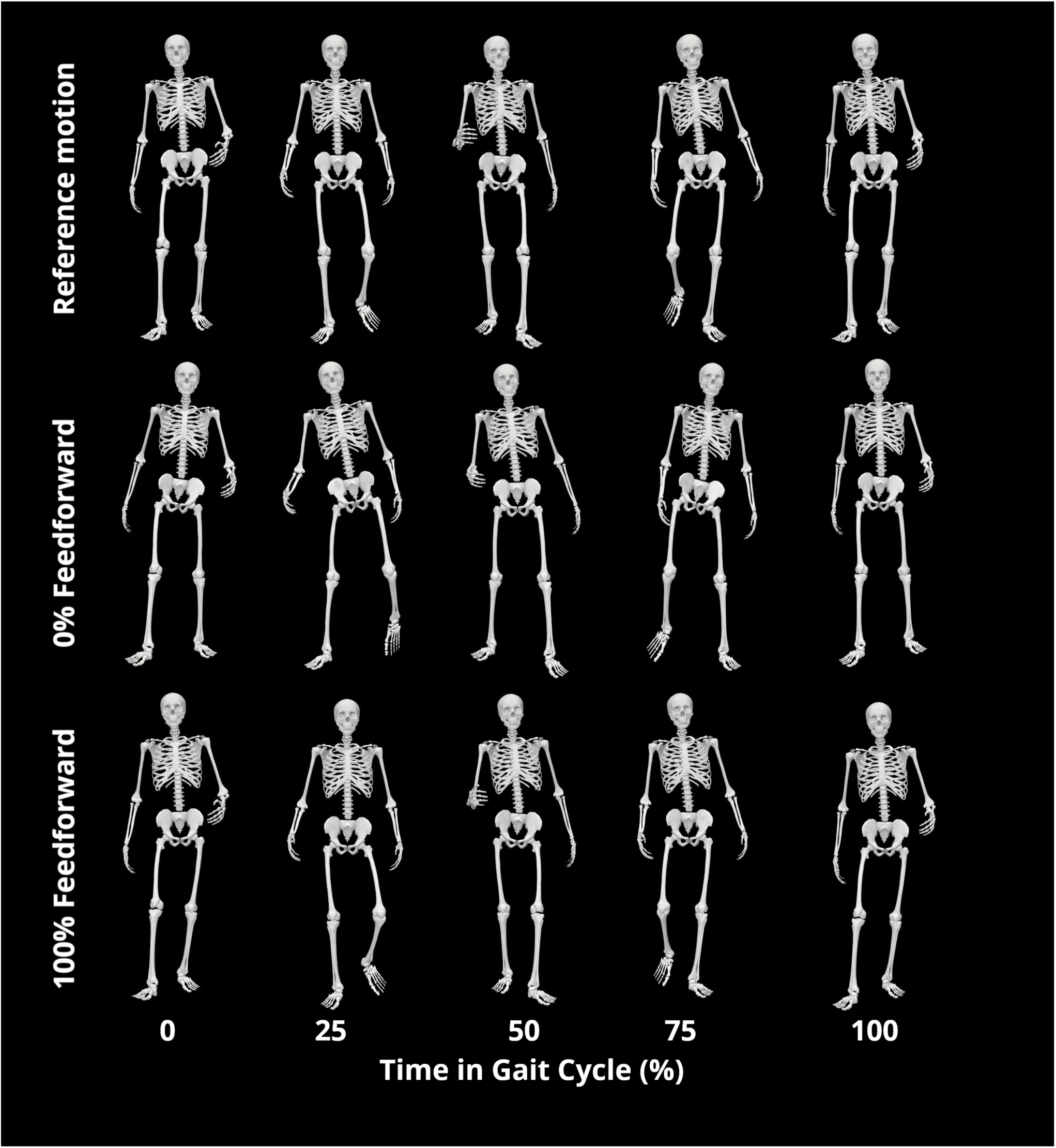
Animation strip of cycle 3 reference motion, motion predicted by feedback model at 0% feedforward contribution without initial value constraints, and motion predicted by feedback model at 100% feedforward contribution.

**Table 2.** Root mean square error (RMSE) (mean ± standard deviation) in predictions of joint angles included on right (R) and left (L) sides as feedback model inputs for each FF contribution level and all 3 testing gait cycles. For comparison, torque-based results are included for the 3 testing gait cycles. Errors are reported in degrees.

| FF contribution | 0% | 25% | 50% | 75% | 100% | 125% |
| --- | --- | --- | --- | --- | --- | --- |
| Hip flex. (R) | 16.3 ± 0.93 | 12.8 ± 0.72 | 8.45 ± 1.27 | 7.06 ± 0.79 | 2.89 ± 1.71 | 2.41 ± 1.15 |
| Hip add. (R) | 12.3 ± 3.01 | 8.83 ± 1.49 | 4.93 ± 0.40 | 1.84 ± 0.50 | 2.01 ± 0.09 | 2.55 ± 0.41 |
| Knee angle (R) | 19.7 ± 3.00 | 17.9 ± 0.21 | 15.3 ± 0.58 | 11.0 ± 0.90 | 4.87 ± 0.67 | 6.49 ± 0.60 |
| Ankle angle (R) | 12.5 ± 0.50 | 8.60 ± 0.90 | 9.56 ± 1.08 | 3.52 ± 0.26 | 1.45 ± 0.20 | 2.93 ± 0.95 |
| Subtalar (R) | 14.5 ± 1.53 | 5.35 ± 0.68 | 12.3 ± 2.10 | 4.05 ± 0.64 | 5.29 ± 0.43 | 3.61 ± 0.29 |
| Hip flex. (L) | 15.7 ± 1.80 | 9.98 ± 0.88 | 6.44 ± 0.93 | 4.36 ± 1.59 | 4.36 ± 0.71 | 6.10 ± 0.57 |
| Hip add. (L) | 13.4 ± 2.76 | 9.85 ± 1.27 | 5.86 ± 0.40 | 2.87 ± 0.71 | 2.51 ± 0.99 | 4.73 ± 1.09 |
| Knee angle (L) | 15.4 ± 1.02 | 13.3 ± 1.11 | 12.6 ± 1.71 | 7.66 ± 1.20 | 6.03 ± 0.37 | 9.25 ± 1.25 |
| Ankle angle (L) | 9.71 ± 1.36 | 7.54 ± 1.19 | 10.9 ± 1.11 | 4.48 ± 0.84 | 2.12 ± 0.44 | 2.64 ± 0.08 |
| Subtalar (L) | 16.7 ± 3.43 | 8.38 ± 0.87 | 21.7 ± 2.29 | 8.86 ± 1.23 | 7.72 ± 0.78 | 5.87 ± 1.74 |

**Table 3.** RMSE (mean ± standard deviation) in predictions of joint moments included as feedback model inputs for each FF contribution level and all 3 testing gait cycles. For comparison, torque-based results are included for the 3 testing gait cycles. Errors are reported in Nm.

| FF contribution | 0% | 25% | 50% | 75% | 100% | 125% |
| --- | --- | --- | --- | --- | --- | --- |
| Hip flex. (R) | 15.9 ± 2.35 | 11.1 ± 2.17 | 6.17 ± 1.69 | 4.23 ± 0.73 | 5.35 ± 1.58 | 6.85 ± 1.38 |
| Hip add. (R) | 33.7 ± 0.88 | 25.6 ± 0.77 | 14.8 ± 1.04 | 7.19 ± 0.60 | 3.41 ± 0.40 | 8.43 ± 1.01 |
| Knee angle (R) | 13.3 ± 2.53 | 15.4 ± 2.67 | 16.9 ± 3.29 | 12.9 ± 1.85 | 5.61 ± 2.10 | 8.42 ± 1.59 |
| Ankle angle (R) | 27.9 ± 3.10 | 21.5 ± 2.25 | 15.5 ± 2.34 | 8.41 ± 2.17 | 9.00 ± 3.01 | 21.2 ± 2.82 |
| Subtalar (R) | 2.82 ± 0.97 | 1.44 ± 0.79 | 3.03 ± 1.02 | 1.67 ± 0.47 | 1.43 ± 0.46 | 2.74 ± 0.25 |
| Hip flex. (L) | 20.2 ± 1.23 | 15.6 ± 1.46 | 10.6 ± 1.09 | 6.06 ± 1.96 | 5.85 ± 2.47 | 8.02 ± 3.23 |
| Hip add. (L) | 41.6 ± 1.46 | 34.9 ± 0.76 | 23.6 ± 1.27 | 11.9 ± 0.52 | 5.98 ± 1.05 | 8.68 ± 2.68 |
| Knee angle (L) | 8.26 ± 1.52 | 8.53 ± 2.18 | 9.37 ± 2.54 | 7.79 ± 2.58 | 7.49 ± 2.38 | 7.64 ± 1.59 |
| Ankle angle (L) | 26.4 ± 7.16 | 23.8 ± 6.40 | 18.6 ± 6.79 | 11.4 ± 6.95 | 9.64 ± 1.96 | 11.8 ± 1.29 |
| Subtalar (L) | 3.24 ± 0.84 | 3.17 ± 0.83 | 4.38 ± 0.41 | 2.16 ± 0.34 | 1.76 ± 0.44 | 1.90 ± 0.18 |

**Table 4.** RMSE (mean ± standard deviation) in predictions of ground reactions (anterior force, vertical force, lateral force, frontal moment, transverse moment, and sagittal moment) for each FF contribution level and all 3 testing gait cycles. For comparison, torque-based results are included for the 3 testing gait cycles. Errors are reported in N for forces and Nm for moments.

| FF contribution | 0% | 25% | 50% | 75% | 100% | 125% |
| --- | --- | --- | --- | --- | --- | --- |
| Ant. F (R) | 38.6 ± 2.23 | 29.0 ± 4.74 | 19.9 ± 3.72 | 13.3 ± 1.93 | 13.5 ± 2.84 | 22.4 ± 6.75 |
| Vert. F (R) | 172 ± 35.3 | 128 ± 11.3 | 63.4 ± 9.94 | 58.4 ± 3.10 | 59.5 ± 24.5 | 70.2 ± 20.5 |
| Lat. F (R) | 27.6 ± 2.30 | 21.5 ± 3.85 | 15.0 ± 4.87 | 11.2 ± 4.79 | 9.71 ± 2.31 | 9.39 ± 1.82 |
| Front. M (R) | 5.17 ± 0.38 | 2.62 ± 0.59 | 2.65 ± 1.10 | 1.83 ± 0.32 | 1.54 ± 0.46 | 2.77 ± 0.32 |
| Trans. M (R) | 4.54 ± 0.28 | 3.17 ± 0.39 | 2.00 ± 0.17 | 1.85 ± 0.21 | 1.89 ± 0.20 | 3.36 ± 0.54 |
| Sag. M (R) | 26.4 ± 2.46 | 23.0 ± 1.99 | 16.0 ± 2.70 | 9.15 ± 2.07 | 7.54 ± 2.99 | 20.6 ± 3.00 |
| Ant. F (L) | 44.2 ± 1.41 | 29.4 ± 2.86 | 12.7 ± 3.26 | 16.1 ± 3.62 | 25.7 ± 6.30 | 23.1 ± 4.88 |
| Vert. F (L) | 165 ± 18.5 | 115 ± 26.0 | 67.1 ± 10.5 | 62.1 ± 8.08 | 60.9 ± 13.2 | 70.2 ± 17.8 |
| Lat. F (L) | 31.9 ± 5.81 | 23.4 ± 0.42 | 19.8 ± 1.06 | 19.1 ± 0.45 | 11.8 ± 0.97 | 11.0 ± 2.92 |
| Front. M (L) | 5.45 ± 1.26 | 3.83 ± 0.57 | 3.69 ± 0.64 | 2.18 ± 1.05 | 2.41 ± 0.26 | 2.34 ± 0.44 |
| Trans. M (L) | 4.31 ± 0.25 | 2.07 ± 0.36 | 1.76 ± 0.29 | 2.94 ± 0.84 | 1.18 ± 0.20 | 2.31 ± 0.42 |
| Sag. M (L) | 28.9 ± 8.53 | 26.5 ± 6.13 | 15.3 ± 5.99 | 11.6 ± 6.16 | 9.50 ± 1.79 | 11.6 ± 1.80 |

## 4 Discussion

This study developed a subject-specific neuromusculoskeletal model with a synergy-based FF+FB lower body control model capable of reproducing walking motions measured from an actual human subject. To our knowledge, this study is the first to perform predictive simulations of walking using a combined FF+FB control model applied to a personalized 3D neuromusculoskeletal model and to evaluate the predictions against actual walking data collected from the subject being modeled. As the simplest possible starting point, the study used FB models with joint-level (rather than muscle-level) inputs and synergy-level (rather than muscle-level) outputs. For the three testing gait cycles, a rather significant level of nominal synergy FF control (at least 75%) was needed for combined synergy-based FF+FB control to succeed in predicting near-periodic walking motions that remained close to experimental initial conditions. While lower levels of nominal FF control were unsuccessful at generating such walking motions, they were able to predict near-periodic walking motions that deviated substantially from the subject’s experimental motion, suggesting a potentially important role of feedforward control for real-life walking. Our results indicate that while fully feedback-controlled simple models may walk in predictive simulations (Geyer and Herr, 2010), researchers should be cautious about applying such models to actual human subjects or when seeking to replicate actual 3D experimental walking data.

This study faced multiple technical challenges that needed to be overcome before synergy-based FF+FB control models could be constructed from actual experimental walking data. First, an appropriate number of synergies to use for the neural control structure needed to be determined. We use of 6 synergies per leg produced a slightly higher total VAF than the 5-synergy solution, a neural feedback model developed using 6 synergies was not as successful as the one developed using 5 synergies. On closer inspection of the 6-synergy solution, we found that the synergy vector weights were poorly distributed, with two synergy vectors representing highly similar coordination patterns. This redundancy may be a poor match for a neural feedback model that needs to generalize to predict motions not included in fitting data. Second, the optimization used to fit feedback gains required a regularization term that penalized high individual feedback gains. While the solution for feedback gains was theoretically unique, it was practically non-unique, resulting in high feedback gains when the regularization term was omitted. Adding a small amount of regularization had little effect on fitting accuracy while simultaneously allowing predictive simulations with combined FF+FB to became successful. Third, development of successful neural feedback models required fitting data from multiple gait cycles. As an initial test of our neural feedback models, we fitted a FB model to a single calibration gait cycle using the same nominal FF synergy activations calculated from the five calibration gait cycles. When we then performed a predictive simulation for that single calibration gait cycle, we expected the simulation to work better than any of our predictions for testing cycles, since the FB model produced a near-perfect fit to the synergy activations produced by the NCP tool. However, this model did not converge to a perfect walking solution, generating worse predictions than the 100% FF results for testing cycles. The FB model struggled to make the small changes in control needed to achieve dynamic consistency, since the synergy activations used for FB model fitting did not come from a dynamically consistent simulation but rather from the NCP tool. Thus, it appears that fitting a broader range of data less accurately is more important than fitting a small subset of data almost perfectly, even when predicting a walking motion that is highly consistent with the calibration data. Future research should assess how variability in training data affects the predictive ability of neural feedback models when simulating real-life walking data.

Since this study investigated synergy-based FF+FB control of walking, an interesting question is whether FB control was needed at all for the three testing cycles. To explore this question, we performed additional predictive simulations for the three testing walking cycle using 75% and 100% FF control with no FB whatsoever. Somewhat to our surprise, the 100% FF case with no FB was successful for all three testing cycles. In contrast, the 75% FF case was successful for only one of the three testing cycles, and even then, the predicted walking motion exhibited almost no knee flexion during stance phase, presumably to reduce the required activation of knee extensor muscles (see Supplementary Material Tables 9 through 11for further details). These observations are consistent with finding that significant FF control (greater than 75%) with some amount of FB is needed to produce experimentally measured walking motions.

These additional simulations highlight an important distinction between the way we scaled our nominal FF synergy activations and the percentage of control that can be categorized as FF. In the two predictive simulations without FB, the percentage of FF control was 100% by definition, since FB made no contribution to the control. Similarly, in our predictive simulations using 100% scaled FF control with FB, FF was not providing 100% of the control. In our synergy-based control models, FF controls are always positive, while FB controls can be positive or negative. Consequently, there is no unambiguous way to calculate the percentage of FB control for this situation. We therefore propose to quantify “percentage FB control” by taking either the root mean square (RMS) value or maximum absolute value of FB and FF control and calculating

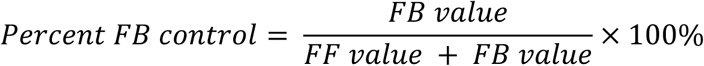

Using this approach, our 75%, 100%, and 125% FF simulations correspond to 4.9 ± 0.1%, 2.2 ± 0.1%, and 3.2 ± 0.2%, respectively using RMS values and 7.3 ± 1.5%, 3.5 ± 1.0%, and 4.5 ± 0.3%, respectively, using maximum absolute values. Either way, the amount of FB control is small relative to the amount of FF control for these successful predictive simulations, and the approach we used for calculating how much nominal FF control was used can be misleading.

Several previous studies have used feedback control to simulate walking (Geyer and Herr, 2010; Jansen et al., 2014; Falisse et al., 2018; Haeufle et al., 2018), but the present study is unique in its modeling fidelity, use of actual human subject walking data, and predictive ability for testing actual walking data. Many feedback models have been two-dimensional (Geyer and Herr, 2010; Haeufle et al., 2018; Aoi et al., 2019, p. 20; Di Russo et al., 2021; Lassmann et al., 2023; Su and Gutierrez-Farewik, 2023; Koseki et al., 2024), which would fail to capture important movement patterns such increased stance width. Models are typically not personalized to subject movement data (Geyer and Herr, 2010; Song and Geyer, 2013; Haeufle et al., 2018; Aoi et al., 2019; Di Russo et al., 2021; Lassmann et al., 2023; Su and Gutierrez-Farewik, 2023, p. 202; Koseki et al., 2024), limiting the ability of these models to represent a specific subject (Gaudio et al., 2023). Walking motions predicted with feedback control are often compared to experimental data only in broad, population-level terms (Song and Geyer, 2013; Di Russo et al., 2021; Lassmann et al., 2023; Su and Gutierrez-Farewik, 2023; Koseki et al., 2024). While experimental EMG data have been used to fit feedback models (Falisse et al., 2018), this study used EMG data to fit feedback models that, when used in predictive simulations, made the model of the subject walk consistent with the subject’s experimental walking data. These advances were made possible by the model personalization and predictive simulation capabilities built into the NMSM Pipeline software.

An important question for this study is whether the source (joint-instead of muscle level) and target (synergy-instead of muscle-level) of our sensory FB model is physiologically justifiable. Neuromechanical models of sensory FB typically use muscle-level quantities (e.g., muscle lengths and velocities for stretch reflexes, tendon forces for Golgi tendon organ reflexes) as inputs to sensory FB models, with sensory FB signals contributing to individual muscle controls. In contrast, to simplify our neural FB models, we used joint-level quantities (i.e., joint angles, joint velocities, joint moments) as surrogate inputs for muscle-level quantities in our sensory FB models, with our sensory FB signals contributing to synergy controls. We chose this simplified approach for two reasons. First, FB models at the muscle-level require calibration of a much larger number of FB gains. Since we calibrated our FB models using data from only five walking cycles performed at a single speed, we would have created an overfitting problem by modeling FB at the level of individual muscles. Second, combined FF+FB control models have not been previously investigated using a 3D personalized neuromusculoskeletal walking model that closely reproduced experimental walking data collected from the subject being modeled. Thus, for a first investigation using real human subject data, we chose to start with the simplest representation of neural FB – FB from joint-level quantities to synergy-level controls. Now that the present study has demonstrated combined FF+FB control can work for this simple FB case using experimental walking data and a personalized 3D neuromusculoskeletal walking model, a future study should take the next step by exploring muscle-level FB models using a realistic walking model and real subject experimental walking data. Although this study presents significant advances in modeling neural feedback for a real subject, this work has several important limitations that future research should address. First, a subset of joint angles is not controlled by the feedback model, most notably hip internal rotation and toes rotation. Thus, muscles do not provide complete control of all DOFs in the legs. We attempted to control hip internal rotation with our synergy-based FF+FB control models, but these simulations produced unrealistic joint angles – an issue that was largely absent when this coordinate was removed. A previous Muscle-tendon Model Personalization effort using the same experimental dataset starting from the same generic full body OpenSim model also had difficulty matching the hip internal-external rotation moments, likely due to inaccurate moment arms or peak isometric strength values for hip internal-external rotation muscles(Meyer et al., 2016). Second, no muscle-specific feedback was modeled, even though muscle-specific feedback is important for feedback mechanisms such as the stretch reflex and Golgi tendon reflex (Geyer and Herr, 2010; Jansen et al., 2014; Trompetto et al., 2014; Ang et al., 2018; Falisse et al., 2018; Haeufle et al., 2018). Accurately modeling muscle-specific feedback becomes more complex in detailed 3D models as relationships between agonist-antagonist muscle pairs are not always well-defined and can vary depending on the state of the model or the coordinate of interest. Identifying how to include these interactions in a feedback model with detailed 3D geometry is significant and important work. Using muscle-specific feedback inputs and outputs would also greatly increase the number of feedback gains that must be fitted, creating a more complex optimization problem that could easily suffer from overfitting. Feedback from and to individual muscles could give our model finer control and improve its predictive ability, which would be worthwhile area for future study. Fifth, our muscle models are coupled to rigid tendons and do not account for short-range stiffness. Rigid tendons are likely sufficient for walking at slow speeds (Millard et al., 2013; De Groote et al., 2016), but short-range stiffness is important for even slight movements (De Groote et al., 2017). Fifth, our feedback model is instantaneous and does not include a time delay between feedback input and output quantities. This delay is a known component of feedback models (Geyer and Herr, 2010), though similar past work with complex neuromusculoskeletal models has not included it (Jansen et al., 2014). This moission was due to a technical limitation of the TO tool. The MTP tool applies a variable time delay to EMG data, but as the EMG data are not changed during the optimization, it only needs to evaluate splines fit to the EMG data at specified time points. In the TO tool, our feedback model inputs of joint angles, joint velocities, and joint moments would change with each iteration of the optimization, so a similar approach would require fitting new splines to data at each iteration, which would be a much more complex and computationally expensive operation. Sixth, ground reaction quantities were not included in our feedback model inputs. These quantities are important to the timing of feedback mechanisms (Geyer and Herr, 2010), but their high sensitivity to small kinematic changes made them difficult to include as a feedback model input during optimal control simulations. For this reason, we used joint moments as a surrogate load quantity. Seventh, loads generated by synergy controls do not perfectly match ID joint loads, and the walking models were not perfectly dynamically consistent. In FF+FB controlled simulations, the dynamic consistency constraints described in our Methods allow for discrepancies of up to 1 Nm for matching synergy-generated jont moments to ID joint moments as well as 1 N and 0.1 Nm for pelvis residual loads, respectively. Setting bounds for these constraints that are too tight will cause simulations to fail to converge, but looser constraints allow for less dynamically consistent solutions.

In conclusion, this study presented a novel method of producing predictive gait simulations that use combined synergy-based FF+FB control and can closely reproduce real 3D experimental walking data. The method utilized the NMSM Pipeline to incorporates dense, subject-specific data for highly personalized models and solutions, bringing new possibilities to OpenSim modeling familiar to researchers. With further work in this area using additional walking data collected from additional subjects, it may eventually become possible to fit accurate neural feedback models to walking data collected from individuals with neurological disorders and then use these models in predictive simulations to design improved patient-specific treatments for walking impairments.

## 5 Conflict of Interest

The authors declare that the research was conducted in the absence of any commercial or financial relationships that could be construed as a potential conflict of interest.

## 6 Author Contributions

SW performed all neuromusculoskeletal modeling work after Joint Model Personalization, formulated and ran all other NMSM Pipeline tools, developed code for fitting feedback models and using them with the Tracking Optimization tool, and participated in writing the manuscript draft. GL performed initial neuromusculoskeletal modeling, including model scaling and Joint Model Personalization, and processed all data used in this study. BF supervised this project, assisted with optimal control problem formulations, evaluated optimal control results, and helped with revising the manuscript draft.

## 7 Funding

This study was funded by the National Institutes of Health under grant R01 EB030520 and by the National Science Foundation under grant 1842494.

## 8 Acknowledgements

This article was published as a preprint on bioRxiv (Williams et al., 2026).

## 10 Data Availability Statement

Experimental data, results, and code used to generate the results of this study are available here on SimTK.org.

## Notes

https://simtk.org/projects/synergyfeedback

### Competing Interest Statement

The authors have declared no competing interest.

### Summary of Updates

Final accepted manuscript after first revision in response to reviewer comments.

